# Discovery and biosynthesis of the antibiotic bicyclomycin in distant bacterial classes

**DOI:** 10.1101/236828

**Authors:** Natalia M. Vior, Rodney Lacret, Govind Chandra, Siobhán Dorai-Raj, Martin Trick, Andrew W. Truman

## Abstract

Bicyclomycin (BCM) is a clinically promising antibiotic that is biosynthesised by *Streptomyces cinnamoneus* DSM 41675. BCM is structurally characterized by a core cyclo(L-Ile-L-Leu) 2,5-diketopiperazine (DKP) that is extensively oxidized. Here, we identify the BCM biosynthetic gene cluster, which shows that the core of BCM is biosynthesised by a cyclodipeptide synthase and the oxidative modifications are introduced by five 2-oxoglutarate-dependent dioxygenases and one cytochrome P450 monooxygenase. The discovery of the gene cluster enabled the identification of BCM pathways encoded in the genomes of hundreds of *Pseudomonas aeruginosa* isolates distributed globally, and heterologous expression of the pathway from *P. aeruginosa* SCV20265 demonstrated that the product is chemically identical to BCM produced by *S. cinnamoneus*. Overall, putative BCM gene clusters have been found in at least seven genera spanning *Actinobacteria* and *Proteobacteria* (*Alpha-, Beta-* and *Gamma-*). This represents a rare example of horizontal gene transfer of an intact biosynthetic gene cluster across such distantly related bacteria, and we show that these gene clusters are almost always associated with mobile genetic elements.

**IMPORTANCE:** Bicyclomycin is the only natural product antibiotic that selectively inhibits the transcription termination factor Rho. This mechanism of action, combined with its proven biological safety and its activity against clinically relevant Gram-negative bacterial pathogens, makes it a very promising antibiotic candidate. Here, we report the identification of the bicyclomycin biosynthetic gene cluster in the known producing organism *Streptomyces cinnamoneus*, which will enable the engineered production of new bicyclomycin derivatives. The identification of this gene cluster also led to the discovery of hundreds of bicyclomycin pathways encoded in highly diverse bacteria, including the opportunistic pathogen *Pseudomonas aeruginosa*. This wide distribution of a complex biosynthetic pathway is very unusual, and provides an insight into how a pathway for an antibiotic can be transferred between diverse bacteria.

## INTRODUCTION

Bicyclomycin (BCM) is a broad-spectrum antibiotic active against Gram-negative bacteria that was first isolated in 1972 from *Streptomyces cinnamoneus* (originally named *Streptomyces sapporoensis*) (1) and is also produced by two other *Streptomyces* species (2, 3). BCM (also known as bicozamycin) is one of the most complex members of the 2,5-diketopiperazine (DKP) family of molecules, cyclic dipeptides generated by the head-to-tail condensation of two α-amino acids (4). The core DKP of BCM, cyclo(L-Ile-L-Leu) (cIL), is modified with a characteristic second cycle that forms a [4.2.2] bicyclic unit, an exomethylene group and multiple hydroxylations (5) (Fig. 1A). BCM is a selective inhibitor of the transcription termination factor Rho (6), which is an essential protein in many bacteria (7, 8) and has been used to treat traveller’s diarrhea (9), as well as in veterinary medicine to treat calves, pigs and fish (7).

**Figure 1.**
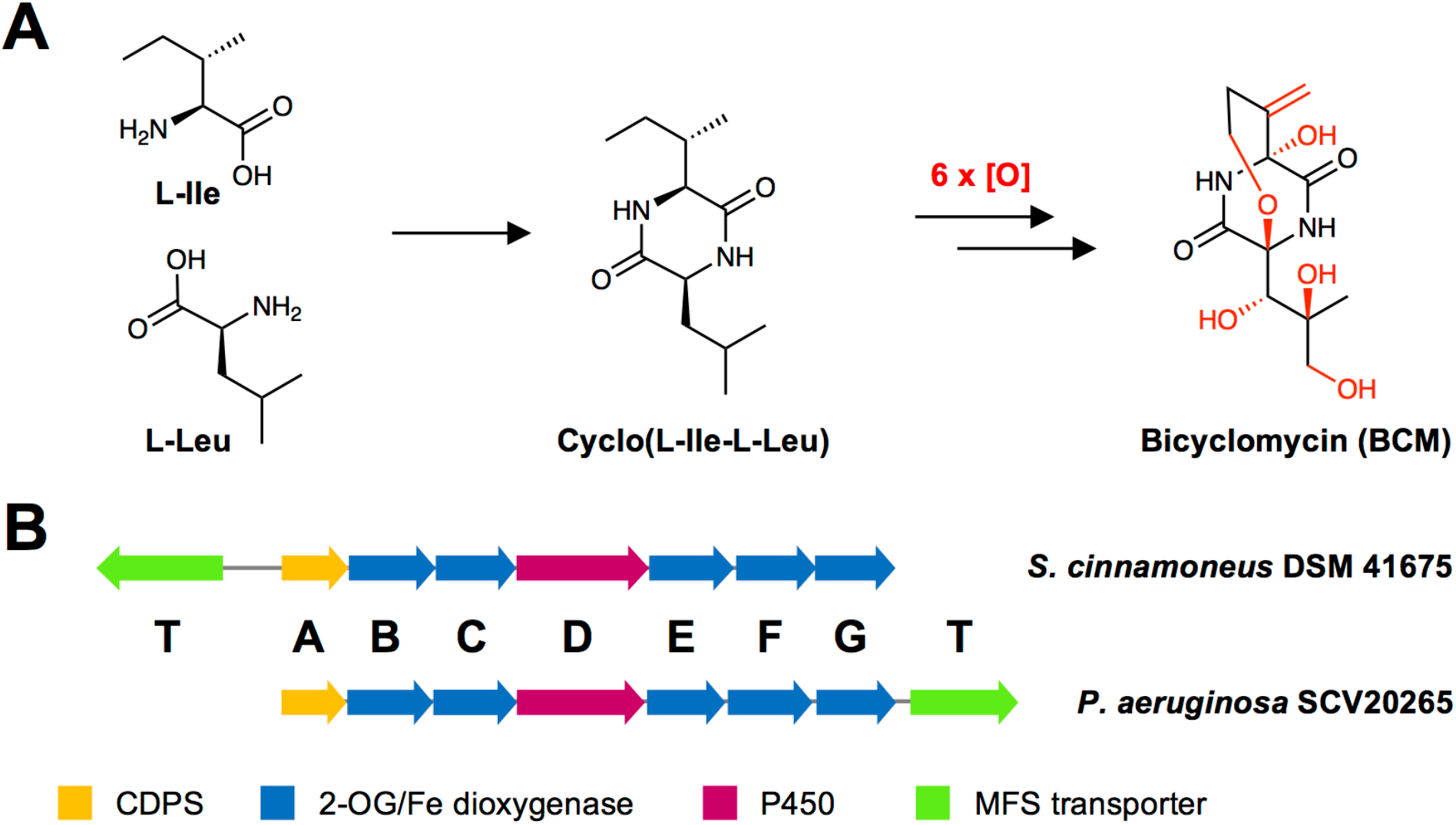
Bicyclomycin biosynthesis. (A) Simplified schematic of BCM biosynthesis. (B) *bcm* gene clusters identified in *S. cinnamoneus* and *P. aeruginosa*.

BCM is the only natural product known to target Rho, which together with its proven safety in mammals and its activity against clinically relevant ESKAPE pathogens like *Acinetobacter baumannii* and *Klebsiella pneumoniae*, makes it a very attractive antibiotic (7, 10). This promise is enhanced by the recent discovery that a combination of BCM with bacteriostatic concentrations of antibiotics targeting protein synthesis leads to a rapid bactericidal synergy (10).

Furthermore, structure-activity relationship studies show that BCM potency can be improved through modification of its exomethylene group (11, 12).

In contrast with the extensive knowledge on BCM mechanism of action (6, 7), very little was known about the biosynthesis of this antibiotic. Feeding experiments previously showed that the DKP scaffold derives from L-leucine and L-isoleucine, as well as the likely involvement of a cytochrome P450 monooxygenase in one of the oxidative steps that convert cIL into BCM (13) (Fig. 1A). To understand BCM biosynthesis, we identified the biosynthetic gene cluster for BCM in *S. cinnamoneus* DSM 41675, which showed that the DKP core is produced by a cyclodipeptide synthase (CDPS) (Fig. 1B). This discovery enabled the identification of homologous clusters in several other species, including hundreds of *Pseudomonas aeruginosa* isolates, an opportunistic pathogen that causes serious hospital-acquired infections. We prove that the *P. aeruginosa bcm* gene cluster is functional and its product is identical to BCM from *Streptomyces*, so represents a viable alternative platform for BCM production. This is a rare example of an almost identical biosynthetic gene cluster in both Gram-negative and Gram-positive bacteria. An analysis of the phylogeny and genomic context of *bcm* gene clusters provides an insight into its likely dispersion through horizontal gene transfer (HGT), and also implies that the *bcm* gene cluster may have undergone a partial genetic rearrangement between Gram-positive and Gram-negative bacteria.

## RESULTS AND DISCUSSION

### Genome sequencing and identification of the BCM gene cluster in *S. cinnamoneus*

The genome sequence of a known BCM producer, *S. cinnamoneus* DSM 41675 was obtained using a combination of Oxford Nanopore MinION and Illumina MiSeq technologies. Illumina MiSeq provided accurate nucleotide level read data, but an Illumina-only assembly was distributed across 415 contigs, in part due to the difficulties in assembling short read data of highly repetitive sequences from large modular polyketide synthase (PKS) and non-ribosomal peptide synthetase (NRPS) genes (14), which were found at the start or end of multiple contigs. Therefore, we also sequenced the genome using Oxford Nanopore MinION technology, which is capable of achieving read lengths of over 150 kb (15), The Nanopore output enabled a much better assembly of the genome over 4 contigs, although at a much lower accuracy at the nucleotide level. Using the raw read data from both sequence runs, we obtained a hybrid assembly composed of a 6.46 Mb contig containing almost all of the chromosome, and a smaller 199 kb contig (Table S1). antiSMASH analysis (16) of this assembly revealed that the 199 kb contig is likely to form part of the chromosome, as the termini of this contig and the 6.46 Mb contig encode different regions of an enduracidin-like gene cluster. In total, these two contigs yield an almost contiguous 6.66 Mb *S. cinnamoneus* genome sequence.

Published feeding experiments indicate that BCM is a DKP derived from L-leucine and L-isoleucine and that a cytochrome P450 is likely to be involved in the pathway (13). Furthermore, a number of additional oxidative reactions are needed to form the final molecule (Fig. 1A). DKPs are produced naturally by either bimodular NRPSs (17, 18) or by CDPSs (19–21) so we expected the biosynthetic gene cluster for BCM to encode either of these enzymatic systems, plus six to seven oxidative enzymes. Analysis of the *S. cinnamoneus* genome sequence with antiSMASH 3.0.5 (16) indicated that there were no suitable NRPS pathways but also no identifiable CDPS pathways. We therefore assessed the genomic regions surrounding every P450 gene in the genome, which revealed the presence of a P450 gene (*bcmD*) that was clustered with genes encoding five 2-oxoglutarate (2OG)-dependent dioxygenases (*bcmB, bcmC, bcmE, bcmF* and *bcmG*), a CPDS gene (*bcmA*) and a gene encoding a major facilitator superfamily (MFS) transporter (*bcmT*) (Fig 1B). Both P450s and 2OG-dependent dioxygenases are capable of catalysing the regiospecific and stereospecific oxidation of non-activated C-H bonds (22–24), while MFS transporters often function as drug-efflux pumps and can confer antibiotic resistance (25, 26).

The putative CDPS (pfam16715) BcmA, has multiple homologs (>45% identity) in other *Actinobacteria* and, notably, in various *Pseudomonas aeruginosa* strains. Interestingly a homolog from *P. aeruginosa* (WP_003158562.1) was previously shown to catalyse the *in vitro* synthesis of cIL (27), and BcmA contains almost all the same specificity-determining binding pocket residues as WP_003158562.1 (Fig. S1). Surprisingly, the five 2OG-dependent dioxygenases encoded in the cluster share only moderate sequence identity (33 to 45%). In total, the gene cluster encodes six oxidative enzymes, which is consistent with the number of modifications required to convert cIL into BCM.

### Heterologous expression of the *bcm* gene cluster

To test whether the identified gene cluster was indeed responsible and sufficient for the biosynthesis of BCM, a 7 kb region spanning *bcmA* to *bcmG* was PCR amplified and cloned into the ϕBT1 integrative vector pIJ10257 (28) by Gibson assembly (29) to generate pIJ-BCM. This places the constitutive promoter ermE*p before *bcmA*, which we anticipated would promote the expression of all *bcm* genes as they are tightly clustered on the same strand. The putative transporter gene *bcmT* was not included on the basis that several homologs of this gene, as well as a homolog of the reported BCM resistance gene (30), are present in the *S. coelicolor* genome. pIJ-BCM was introduced into *S. coelicolor* M1146 and M1152 (31) via intergeneric conjugation. LC-MS^2^ analysis of cultures of the resulting strains yielded a peak of *m/z* 285.11 not present in control strains (Fig. 2), which had an identical retention time and MS^2^ fragmentation pattern (*m/z* 211.05, *m/z* 193.2, *m/z* 108.4 and *m/z* 81.9, Fig. S2) to BCM produced by *S. cinnamoneus*, as well as a pure BCM standard, and corresponds to [BCM-H_2_O+H]^+^. This unambiguously confirmed that this was the BCM biosynthetic gene cluster. Our result agrees with recent studies by Patteson *et al*. (32) and Meng *et al*. (33) who, in parallel with our study, have reconstituted *in vitro* the functions of the CDPS and the oxidative steps in the *S. cinnamoneus* pathway.

**Figure 2.**
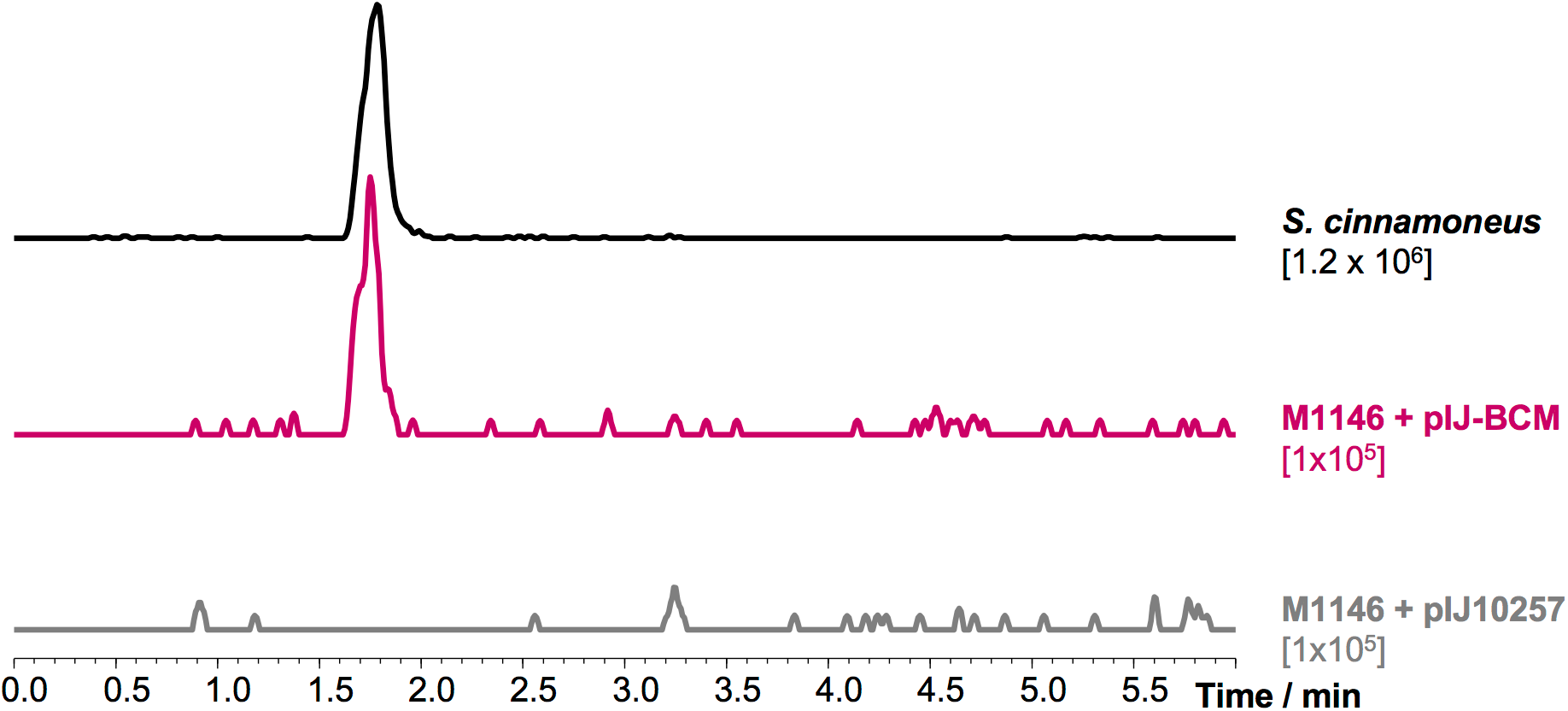
Heterologous expression of the *bcm* gene cluster from *S. cinnamoneus* in *S. coelicolor* M1146. Extracted ion chromatograms (EICs) of bicyclomycin (*m/z* 285.11, [M-H_2_O+H]^+^) in *S. cinnamoneus, S. coelicolor* M1146 expressing the *bcm* cluster and *S. coelicolor* M1146 containing empty vector. The intensity scale of each EIC is noted under the corresponding label.

### Identification and heterologous expression of a *bcm* gene cluster from *Pseudomonas aeruginosa*

During our bioinformatic analysis of the *S. cinnamoneus bcm* gene cluster it became clear that entire bcm-like gene clusters with an apparently identical organisation of *bcmA-G* genes were present a variety of Gram-negative and Gram-positive bacterial species, and in particular in multiple *P. aeruginosa* strains. The widespread distribution of such a conserved antibiotic gene cluster is very rare and prompted us to investigate whether these highly similar gene clusters actually make identical products. As a representative example, *P. aeruginosa* SCV20265 was therefore investigated for its ability to produce BCM. This strain is a well-studied (34–36) small colony variant of the opportunistic pathogen isolated from the lung of a patient with cystic fibrosis (37) and is considered a reference strain in antibiotic resistance studies (38). The *P. aeruginosa* SCV20265 bcm-like gene cluster encodes proteins with sequence identities of between 30–56% compared to their *Streptomyces* counterparts. A MFS transporter is also encoded in this cluster, but is at the end of the *bcmA-G* operon instead of preceding *bcmA* (Fig. 1B).

No BCM production was detected in cultures of *P. aeruginosa* SCV20265, so heterologous expression of the gene cluster was carried out to determine whether the pathway is functional. The putative *bcm* cluster (including *bcmT*) was PCR amplified from SCV20265 gDNA and cloned into pJH10TS (39, 40), which places the putative *bcm* operon under the control of the synthetic promoter Ptac. *Pseudomonas fluorescens* SBW25 was transformed with the resulting plasmid (pJH-BCMclp-PA). Several clones of this heterologous expression strain were cultured in a range of production media, and assessed for their ability to produce BCM. LC-MS^2^ analysis revealed that *P. fluorescens* SBW25-pJH-BCMclp-PA efficiently produces BCM after 14 h of growth (Fig. 3). The putative BCM detected in these samples exhibited the same retention time, mass and fragmentation profile as a pure BCM standard, including MS signals of *m/z* 285.11, as observed previously, and *m/z* 325.10, corresponding to [BCM+Na]^+^ (Fig. 3 and Figs. S3, S4). This result is consistent with parallel work from Patteson *et al*. (32), but this does not preclude the possibility of variation in stereochemistry at one more positions in the molecule. We therefore scaled up production, purified the compound and subjected it to NMR analysis (^1^H, ^13^C, COSY, HMBC, HSQC), which provided identical spectra (Figs. S5 to S10, Table S2) to authentic BCM reported previously (41). *Pseudomonas*-produced BCM also had the same optical rotation as a BCM standard, confirming that they are stereochemically identical.

**Figure 3.**
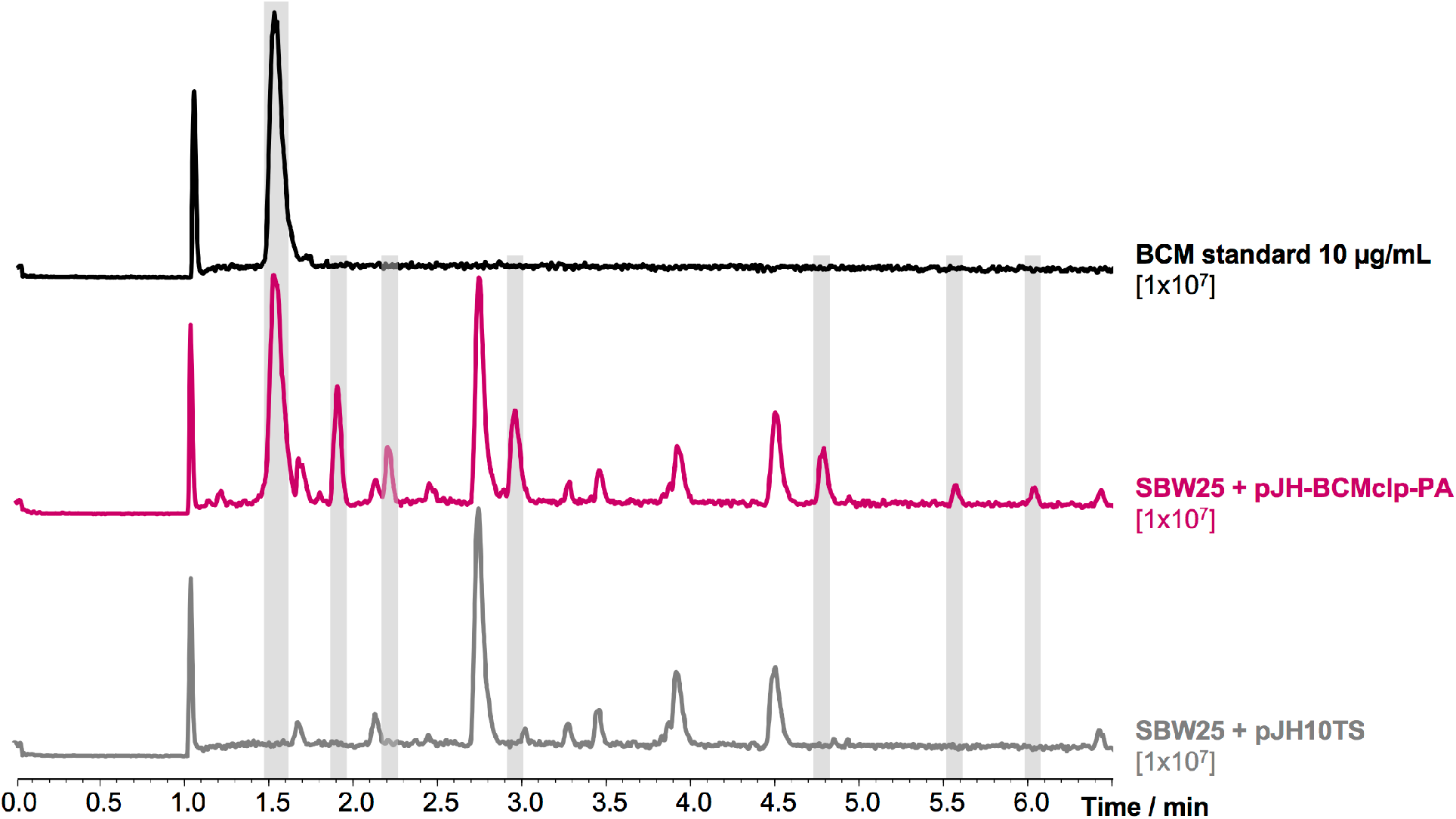
Heterologous expression of the *bcm* cluster from *P. aeruginosa* in *P. fluorescens* SBW25. Base peak chromatograms of a bicyclomycin standard, *P. fluorescens* SBW25 expressing the *bcm* cluster and *P. fluorescens* SBW25 containing empty vector. The intensity scale of the chromatograms is noted under the corresponding labels. Compounds produced by the heterologous expression strain but not found in the control strain are highlighted in grey.

One of the most efficient media for BCM production in *P. fluorescens* was SCFM, a synthetic medium that mimics the salt and amino acid composition from cystic fibrosis sputum samples (42). The composition of this medium was simplified to generate bicyclomycin production medium (BCMM), in which cultures of *P. fluorescens* SBW25-pJH-BCMclp-PA provided BCM yields of 34.5 ± 2.1 mg/L in only 14 h. Interestingly, we could detect at least six additional compounds in the heterologous expression strain in comparison to a negative control strain harbouring empty pJH10TS (Fig. 3 and Figs. S3, S4). All of these compounds have masses compatible with BCM-like compounds (Table S3) and some have BCM-like MS^2^ fragmentation patterns, such as a loss of 74.04 Da that corresponds to fragmentation of the oxidized leucine side chain (Fig. S4). This production profile makes *P. fluorescens* a promising BCM production system when compared to the complex media and longer incubation times required to produce BCM in *Streptomyces* species. In contrast, we could not detect any BCM-like molecules in cultures of wild type *P. aeruginosa* SCV20265, suggesting that additional factors are required to activate the expression of an otherwise functional gene cluster.

### Organisation, taxonomic distribution and phylogeny of the *bcm* cluster

The presence of seven contiguous biosynthetic genes that make the same antibiotic in both Gram-positive and Gram-negative bacteria was a fascinating result. The production of the same compound in such distantly related organisms (bacteria that are evolutionarily at least 1 billion years apart (43)) is incredibly rare, but not unprecedented (44). To investigate this unusual result, a BLASTP search using BcmA was used to identify every putative *bcm* gene cluster (*bcmA-G*) in sequenced bacterial genomes. In total, 724 candidates were identified, where 31 are found in a variety of taxa and the remaining sequences all come from *Pseudomonas* species, in particular *P. aeruginosa*. This initial dataset was filtered (see Material and Methods) to generate a final dataset for phylogenetic analysis containing 374 bcm-like gene clusters (Data set S1). Analysis of this dataset showed that *bcm*-like gene clusters are also found in seven other sequenced *Streptomyces* species besides *S. cinnamoneus*, as well as 20 *Mycobacterium chelonae* strains, *Williamsia herbipolensis* (order *Corynebacteriales), Actinokineospora spheciospongiae* (order *Pseudonocardiales*) and the Gram-negative bacteria *Burkholderia plantarii* and *Tistrella mobilis (Beta-* and *Alphaproteobacteria*, respectively). Furthermore, a fragmented bcm-like gene cluster was identified in *Photorhabdus temperata* (*Gammaproteobacteria*) by BLAST analysis of BcmA and the P450 BcmD. This cluster is split across two different contigs (accession numbers NZ_AYSJ01000007 and NZ_AYSJ01000009), where it is accompanied by transposase genes, and was therefore not included in our dataset.

Most *bcm* gene clusters from Gram-positive bacteria share the same gene organisation, with *bcmT* in a divergent operon upstream of *bcmA*, whereas in all the Gram-negative bacteria (and *Actinokineospora*) *bcmT* is downstream of *bcmG. Streptomyces ossamyceticus* is the only representative that lacks a transporter gene immediately adjacent to the biosynthetic genes. Additionally, the MFS transporters from Gram-positive gene clusters only share 27-30% sequence identity (approx. 40% coverage) with MFS transporters from Gram-negative gene clusters, suggesting that the transporters have been recruited independently in these distant bacteria.

All the *bcm* gene clusters identified in this work were analysed phylogenetically by constructing a maximum likelihood tree from the nucleotide sequence spanning *bcmA-G*. This showed that their evolutionary relationship correlates tightly with the taxonomy of the strains (Fig. 4A). Clusters from Gram-negative (particularly *Pseudomonas*) and Gram-positive bacteria are grouped in completely independent and distant clades, while the clusters from *Burkholderia* and *Tristella* appear at intermediate points between these two groups. Within the Gram-positive clade, the clusters have a higher degree of divergence but are similarly grouped according to the classification of their native species, with the *Williamsia* gene cluster clustering with the *M. chelonae* gene clusters (these two genera belong to the order *Corynebacteriales*) (Fig. 4B). All *P. aeruginosa* gene clusters are ~99% identical to each other (Fig. 4A and Fig. S11), whereas the two most distantly related streptomycete gene clusters share 69% identity and 83% coverage.

**Figure 4.**
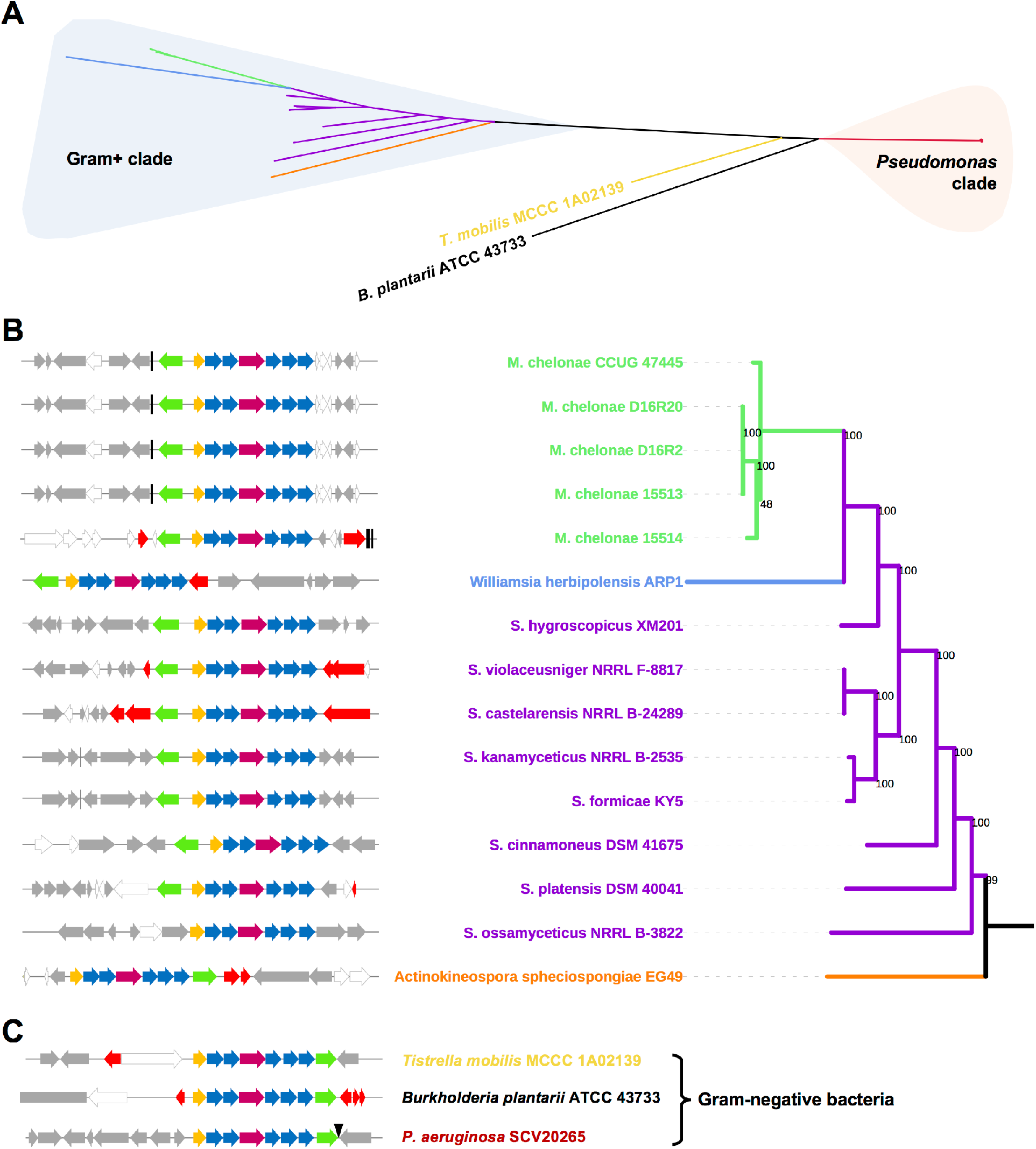
Phylogeny and genetic context of the *bcm* gene clusters. (A) Unrooted maximum likelihood tree of the nucleotide sequences from the *bcm* gene clusters identified in this work. Branches are color-coded by genera and major clades are highlighted. (B) Pruned tree showing gene clusters in Gram-positive bacteria and their genetic context. Bootstrap support values for the phylogeny are shown at the base of each branch and the genetic context of each cluster (color-coded as in Fig. 1B) is shown for each branch of the tree. Flanking genes are color-coded grey if they encode proteins with conserved domains, white for hypothetical proteins with no conserved domains and red for proteins related to mobile genetic elements. Vertical black lines represent tRNAs. (C) Genetic context of the *bcm* clusters in Gram-negative bacteria. The black triangle represents a attTn7 site.

### Mobile genetic elements associated with bcm-like gene clusters

The conserved organisation of biosynthetic genes, along with the phylogenetic relationship between the *S. cinnamoneus* and *P. aeruginosa* CDPSs (32), strongly implies that the *bcm* gene cluster has been horizontally transferred between numerous bacteria. The increased sequence divergence of the *bcm* gene clusters in *Streptomyces* species suggests that the gene cluster may have originated from this taxonomic group, although it is difficult to prove this hypothesis, as the gene clusters in all strains appear to have adapted to their hosts, making HGT difficult to infer. Despite the below average GC content of the clusters (59.6% in *P. aeruginosa* SCV20265 and 70.8% in *S. cinnamoneus*) versus the genome averages (66.3% and 72.4%, respectively), the clusters were not predicted to be part of genomic islands in these strains when analysed with IslandViewer4 (45).

However, analysis of the genomic context of *bcm* gene clusters in *P. aeruginosa* strains strongly supports an insertion hypothesis, since the genes that flank the cluster are contiguous in a number of *P. aeruginosa* strains that lack the cluster (Fig. S12). Most notably, *bcmT* is adjacent to the glucosamine-fructose-6-phosphate aminotransferase gene *glmS*, and the intergenic region that precedes *glmS* contains the specific attachment site for transposon Tn7 (attTn7) (46). Consistent with this observation, some strains that lack the *bcm* gene cluster (e.g. *P. aeruginosa* BL08) have mobile genetic elements integrated next to *glmS* (Fig. S12). Intriguingly, many strains, including the reference strain PAO1, contain a MFS transporter gene (PA5548 in PAO1) adjacent to *glmS* that is 99% identical with *bcmT* from SCV20265. This either indicates that the *bcm* gene cluster recently integrated next to an existing *P. aeruginosa* transporter, or that a subset of strains lost the biosynthetic genes but retained a potential BCM resistance gene.

The bcm-like gene clusters in other Gram-negative bacteria (*Burkholderia* and *Tistrella*) and most Gram-positive bacteria are located next to genes coding for integrases, transposases and other genetic mobility elements, which strongly supports HGT of the cluster into these taxa (Figs. 4B and 4C). For example, the mycobacterial clusters are found close to tRNA genes, and their flanking genes are syntenic in some *M. abscessus* strains, whereas in other *M. abscessus* strains these genes are separated by a cluster of phage-related genes (Fig. 4B and Fig. S13). In the streptomycetes, the clusters are integrated in different genomic locations, where they are also often associated with mobile genetic elements (Fig. 4B). Across all genera, this indicates that *bcm* gene clusters are almost always located at regions of genomic plasticity.

### Diversity and geographical distribution of the *bcm* cluster in *P. aeruginosa*

The high sequence identity of the *bcm* gene cluster across hundreds of *P. aeruginosa* strains (Fig. S11) along with its consistent genomic context (Fig. 4C) led us to question whether this cluster is truly widespread, or only found in a small subset of *P. aeruginosa* strains over-represented in sequence databases. *P. aeruginosa* isolates have been widely sequenced to evaluate pathogen diversity and evolution (38, 47, 48). As a result, large collections of sequenced clinical isolates are available in the databases, potentially constituting a biased dataset that might lead to an overestimation of *bcm* gene cluster abundance and conservation. Most of the sequences in our final *bcm* dataset come from well-characterised isolate collections. Among them, the Kos collection (38) provides a comprehensive survey of *P. aeruginosa* diversity, and the *bcm* gene cluster is present in nearly 20% of isolates sequenced in this collection (74 out of the 390). To assess the phylogenetic diversity of these strains, we plotted the presence of the *bcm* gene cluster onto the Kos collection phylogenetic tree (38). Strikingly, this showed that nearly all of the bcm-positive strains are found in the PAO1 clade (Fig. 5), but that these come from very diverse locations, including the USA, Mexico, Spain, France, Germany, China, Argentina, Brazil, Colombia, Croatia and Israel, among others. This geographic diversity was further augmented by an analysis of all *P. aeruginosa* strains encoding the pathway (Data set S1). We can therefore conclude that the *bcm* gene cluster is distributed globally, but within a phylogenetically distinct subset of *P. aeruginosa* strains. Given this phylogenetic distribution, it is surprising to note that a *bcmT* gene is also found next to *glmS* in *P. aeruginosa* PA14 (Fig. S12).

**Figure 5.**
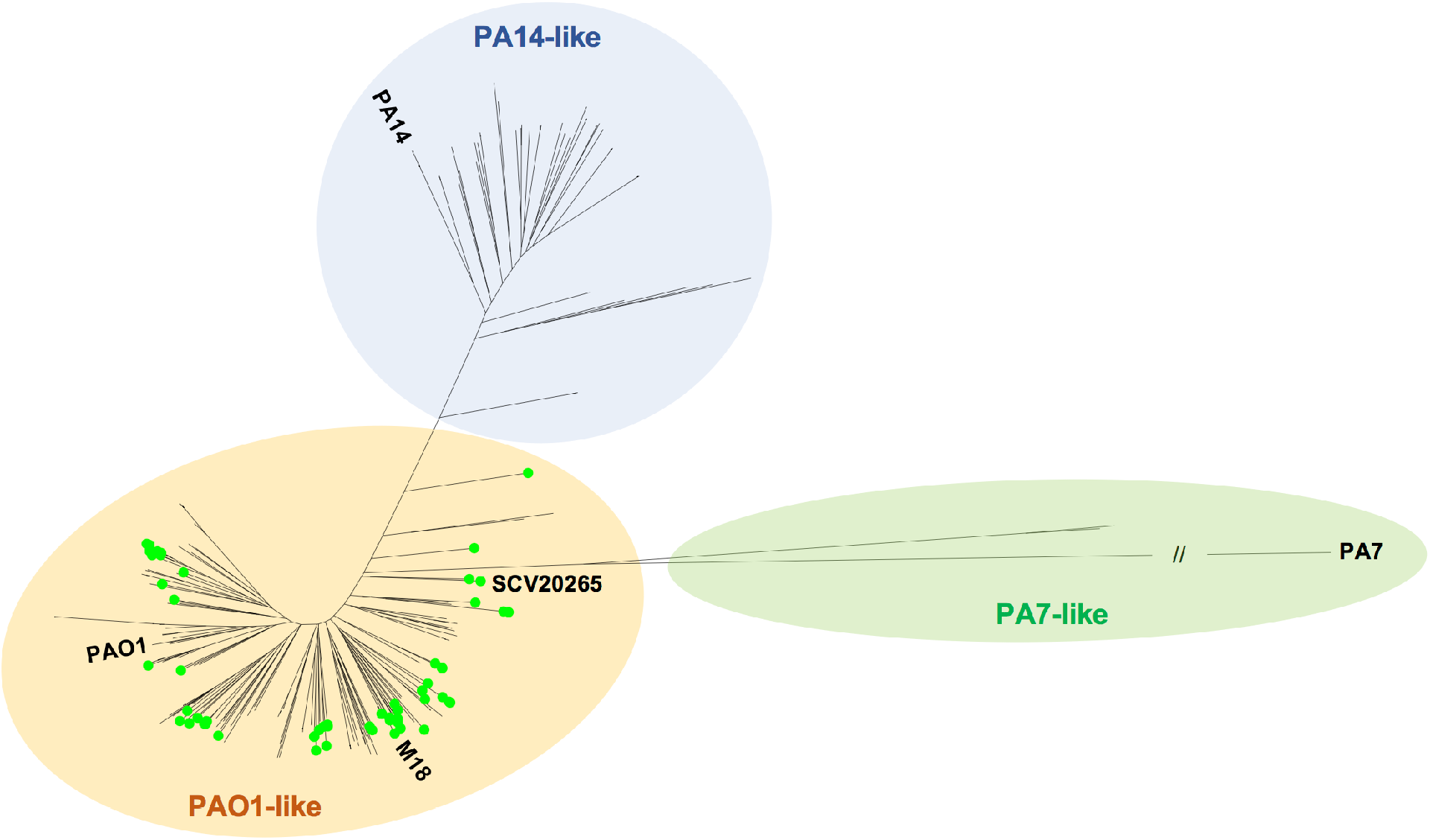
Distribution of the *bcm* gene cluster across *P. aeruginosa* isolates using a modified version of the unrooted maximum-likelihood tree generated by Kos *et al*. (38). The PA14-like, PA7-like and PAO1-like clades are color-coded, and a green dot signifies the presence of the *bcm* gene cluster. *P. aeruginosa* SCV20265 and multiple reference strains (PAO1, M18, PA7, PA14) are also labelled.

### 2OG-dependent dioxygenase phylogeny

An unusual feature of the *bcm* gene clusters is the presence of five 2OG-dependent dioxygenase genes. While it is possible that they originally arose by gene duplication events, the *S. cinnamoneus* 2OG-dependent dioxygenases only possess 33-45% sequence identity with each other (Figure S14). We hypothesised that an analysis of the diversity of the *bcm* 2OG-dependent dioxygenases across multiple taxa could provide an insight into gene cluster evolution. We therefore constructed a maximum likelihood tree using protein sequences of every 2OG-dependent dioxygenase (BcmB, C, E, F and G homologs) from both *S. cinnamoneus* and *P. aeruginosa* SCV20265, as well as from other selected *P. aeruginosa* strains and at least one representative from the other genera that encode bcm-like gene clusters.

In contrast to the overall gene cluster phylogeny, the *bcm* oxidases group primarily based on their position in the cluster, and therefore their likely biosynthetic role (Fig. 6). BcmB, BcmC and BcmG group clearly in different clades, and within these clades the proteins from Gram-negative bacteria branch out from the Gram-positive subgroups, perhaps indicating the ancestral origin of these proteins. A surprising result was the unexpected phylogeny of the remaining two 2OG-dependent dioxygenases, BcmE and BcmF. These are clearly separated into two different clades: one containing BcmE from Gram-negative bacteria (BcmE-) and BcmF from Gram-positive bacteria (BcmF+) and one where BcmE+ groups with BcmF-. Within these two clades, Gram-positive and Gram-negative representatives are more distinct and bifurcate earlier than in the other clades (Fig. 6). This intriguing result might mean that BcmE and BcmF fulfil inverse roles in Gram-positive and Gram-negative bacteria, and further experiments are necessary to test this hypothesis. The phylogenetic relationship between the 2OG-dependent dioxygenases strongly supports HGT of the cluster between taxa, although the BcmE/BcmF phylogeny indicates that the cluster may have undergone some reorganisation (Fig. 6).

**Figure 6.**
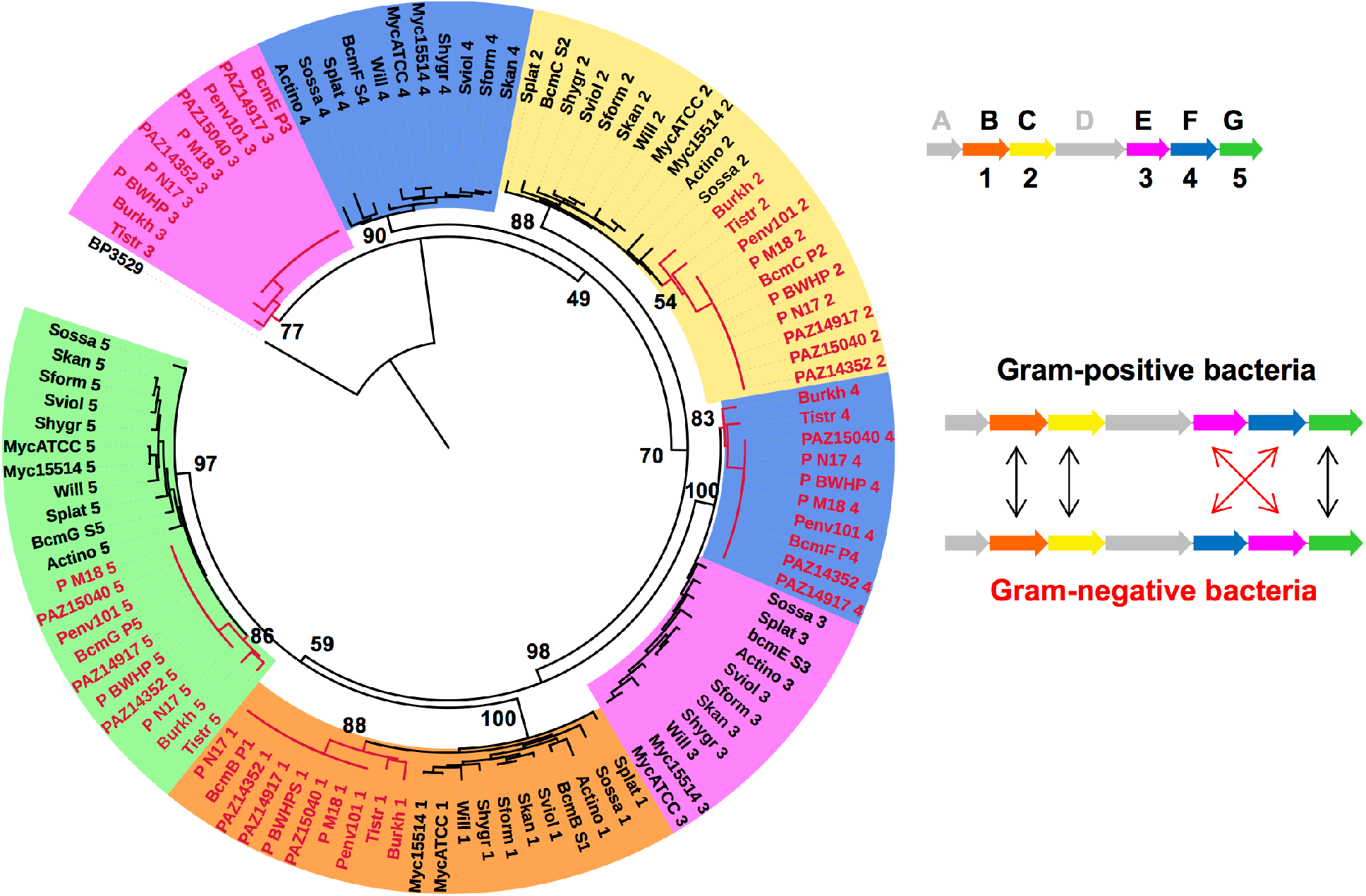
Maximum likelihood tree of the *bcm* 2-OG dioxygenases, including all representatives from *Streptomyces, Actinokineospora, Williamsia, Burkholderia* and *Tistrella*, two from *Mycobacterium*, and eight from *Pseudomonas*. Protein BP3529 from *Bordetella pertussis* was used as outgroup. Background colors and numbering of the branch labels represent the position of a particular dioxygenase in the *bcm* gene cluster, as shown in the schematic representation. The taxonomic origins of each protein are indicated by their branch and label colors (black for Gram-positive and red for Gram-negative representatives). Bootstrap support values for the major branches are shown. A reorganization of the *bcm* gene cluster between Gram-negative and Gram-positive bacteria is proposed based on dioxygenase phylogeny.

In summary, we demonstrate that the antibiotic BCM is a CDPS-derived natural product whose biosynthetic gene cluster is present in a diverse array of both Gram-positive and Gram-negative bacteria. This characterisation was supported by heterologous expression of pathways from *S. cinnamoneus* and *P. aeruginosa*, where the pathway product was proven to be stereochemically identical to authentic BCM. We have also showed that the previously orphan *P. aeruginosa* pathway is a promising system for the production of BCM and related derivatives. The *bcm* cluster is dispersed across a number of taxonomically distant bacteria, including *Alpha-, Beta-* and *Gammaproteobacteria*, as well as several *Actinobacteria* families. The widespread presence of *bcmT* genes in *P. aeruginosa* (even those that lack the biosynthetic genes (Fig. S12)), may explain why BCM is inactive towards *P. aeruginosa* (49), but further work is required to determine whether *bcmT* confers BCM resistance.

The presence of mobile genetic elements associated with the *bcm* gene cluster in many bacteria strongly supports dissemination of this gene cluster via HGT, and the diversity of the gene cluster in Gram-positive bacteria suggests that it then subsequently transferred to Gram-negative bacteria, where two dioxygenase genes have apparently rearranged in the gene cluster and an alternative MFS transporter was acquired. However, the opposite direction of horizontal transfer cannot be ruled out. We are not aware of such a widespread distribution of any other specialized metabolite gene cluster, although there are examples of compounds that have been found in both Gram-positive and Gram-negative bacteria, such as pyochelin (50), the coronafacoyl phytotoxins (51) and furanomycin (52). A recent study by McDonald and Currie showed that It is very rare to find intact laterally transferred biosynthetic gene clusters, even between streptomycetes (53).

Given this distribution of *bcm* gene clusters, it will be interesting to determine the ecological role of BCM, especially given the abundance of functional pathways in pathogenic *P. aeruginosa* strains isolated from lungs, where adaptive evolutionary pressure would have led to the loss or decay of the cluster unless it conferred a competitive advantage (54). Antibacterial natural products can have roles in pathogen virulence, such as a bacteriocin produced by the pathogen *Listeria monocytogenes* that modifies intestinal microbiota to promote infection (55). In addition, given the frequent horizontal transfer of the *bcm* gene cluster and its extensive association with mobile genetic elements, it is interesting to note that transcription terminator Rho most strongly represses transcription of horizontally acquired regions of genomes (56), an activity that would be specifically inhibited by BCM (7). It is known that phages recruit genes from bacteria that increase their fitness and that of their hosts (57, 58) and this may occur with the *bcm* gene cluster. These intriguing observations invite further work to determine the natural role of BCM.

## MATERIAL AND METHODS

### Chemicals and molecular biology reagents

Pharmamedia was obtained from Archer Daniels Midland Company. Antibiotics, and all other media components and reagents were purchased from Sigma-Aldrich. Bicyclomycin was purchased from Bioaustralis Fine Chemicals (Australia). Enzymes were purchased from New England Biolabs unless otherwise specified, and molecular biology kits were purchased from Promega and GE Healthcare.

### Bacterial strains, plasmids and culture conditions

*Escherichia coli, Streptomyces* and *Pseudomonas* strains, as well as plasmids and oligonucleotides used or generated in this work are reported in Tables 1 and 2. *S. cinnamoneus* DSM 41675 was acquired from the German Collection of Microorganisms and Cell Cultures (DSMZ, Germany), *P. aeruginosa* SCV20265 was provided by Prof. Susanne Häussler (Helmholtz Centre for Infection Research, Germany) and pJH10TS was provided by Prof. Barrie Wilkinson (John Innes Centre, UK).

**Table 1.**
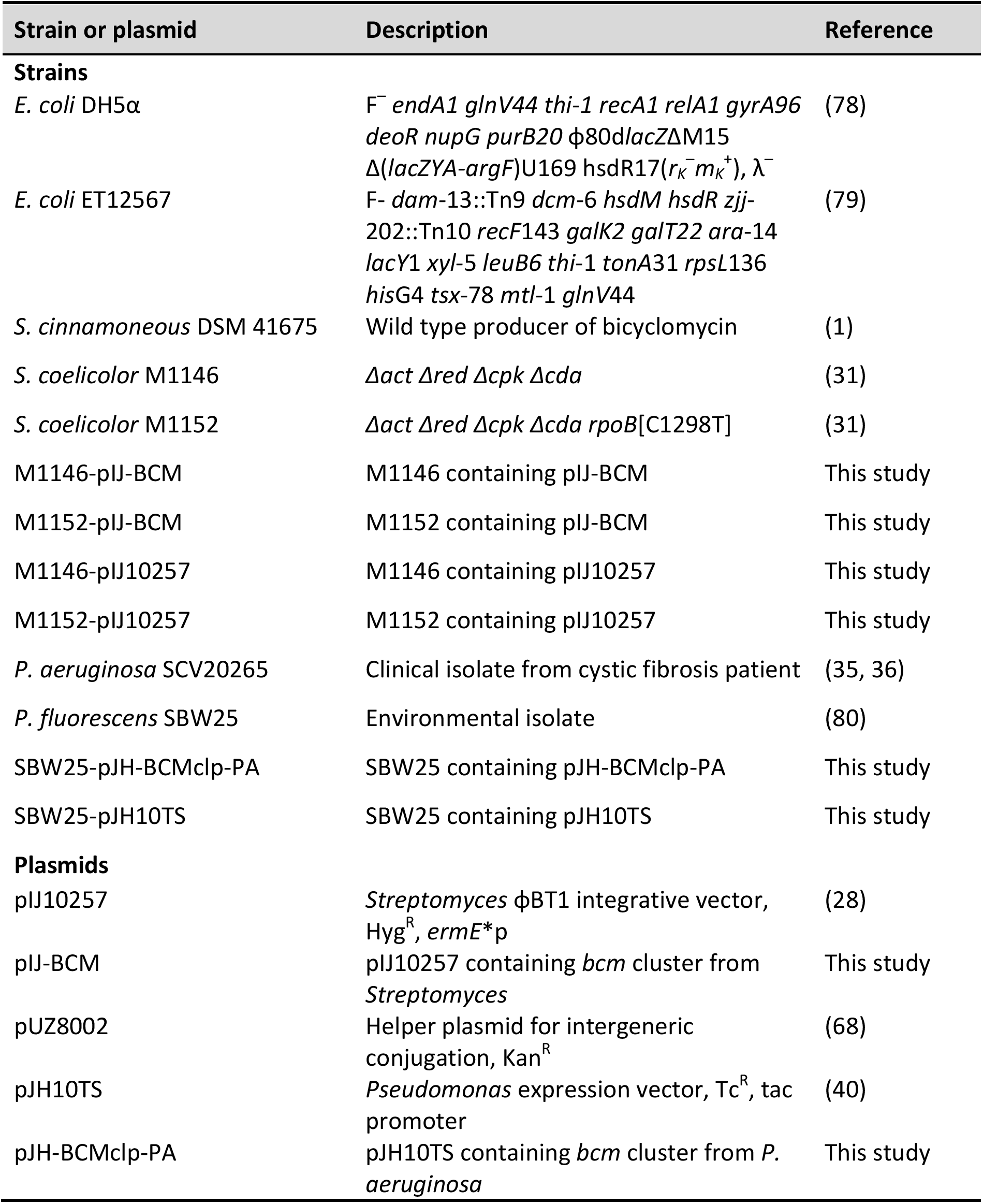
Strains and plasmids used or generated in this study.

**Table 2.**
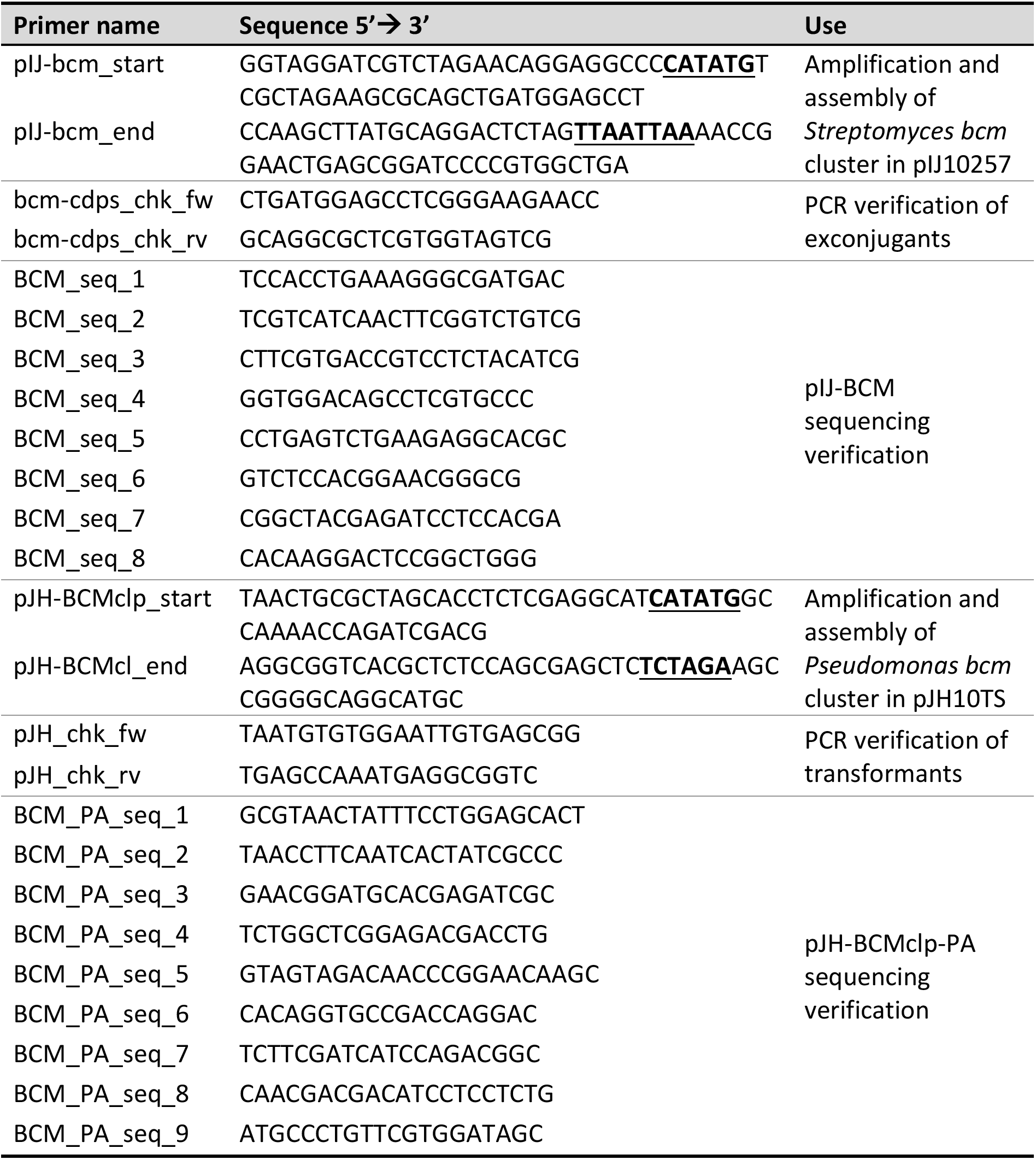
Primers used in this study. Restriction sites used for assembly are underlined and in bold type.

*E. coli* and *Pseudomonas* strains were grown in lysogeny broth (LB) at 37 °C (except for *P. fluorescens* SBW25, which is temperature sensitive and was grown at 28 °C) and stored at −70°C in 50% glycerol stocks. *Streptomyces* strains were cultured in liquid tryptone soya broth (TSB, Oxoid) or solid soya flour mannitol (SFM) medium (59) at 28–30 °C and stored at −70 °C as 20% glycerol spore stocks.

The following media were used for bicyclomycin production experiments: Aizunensis production medium (AIZ), adapted from (60): 20 g/L glucose, 20 g/L soy flour, 2 g/L bactopeptone, 2g/L NaNO_3_, 1 g/L KH_2_PO_4_, 0.5 g/L MgSO_4_·7H_2_O, 0.5 g/L KCl, 0.001 g/L FeSO_4_·7H_2_O, pH 7.0. Cinnamoneus production medium (CIN), adapted from (1): 20 g/L potato starch, 20 g/L cotton seed meal (Pharmamedia), 10 g/L soy flour, 5 g/L MgSO_4_·7H_2_O, 10.9 g/L KH_2_PO_4_, 2.85 Na_2_HPO_4_, pH 6.8 (a solid version of CIN medium with 20 g/L agar was used to grow *S. cinnamoneus* for reliable spore production). Synthetic cystic fibrosis medium (SCFM) was prepared following the recipe reported by Kamath and co-workers (42), and an alternative medium optimized for bicyclomycin production (BCMM) was developed from this (per L): 6.5 mL 0.2 M NaH_2_PO_4_, 6.25 mL 0.2 M Na_2_HPO_4_, 0.348 mL 1 M KNO3, 0.122 g NH_4_Cl, 1.114 g KCl, 3.03 g NaCl, 10 mM MOPS, 16.09 mL 100 mM L-leucine, 11.2 mL 100 mM L-isoleucine, 6.33 mL 100 mM L-methionine, 15.49 mL 100 mM L-glutamic acid hydrochloride, 6.76 mL 100 mM L-ornithine·HCl, 1.92 mL 84 mM L-cystine (dissolved in 0.8 M HCl) and 2 mL 3.6 mM FeSO_4_·7H_2_O, all in milliQ water. The solution was adjusted to pH 6.8, filter sterilised and supplemented with 0.606 mL 1 M MgCl_2_ and 1.754 mL 1 M CaCl_2_ (sterilised separately). When necessary, antibiotics were added at the following concentrations: 50 μg/mL hygromycin, 50 μg/mL apramycin, 50 μg/mL kanamycin, 25 μg/mL chloramphenicol, 25 μg/mL nalidixic acid and 12.5 μg/mL tetracycline.

### Genome sequencing, annotation and bioinformatics analysis of *S. cinnamoneus*

Genomic DNA of *S. cinnamoneus* DSM 41675 was isolated following the salting out protocol (59), which was then subjected to a TruSeq PCR-free library preparation and then sequenced using Illumina MiSeq (600-cycle, 2×300 bp) at the DNA Sequencing Facility, Department of Biochemistry, University of Cambridge (UK). MinION nanopore sequencing (Oxford Nanopore Technologies, UK) was carried out using the following protocol.

A single colony from *S. cinnamoneus* grown on solid CIN medium was used to inoculate 50 mL TSB, which was incubated at 28 °C overnight with shaking at 250 rpm. 1 mL of this seed culture was used to inoculate a further 50 mL TSB, which was again incubated at 28 °C overnight with shaking at 250 rpm. DNA was extracted from 10 mL of this culture using the salting out procedure as described before (59) and resuspended in 5 mL Tris-EDTA (TE) buffer. DNA concentration was quantified using a Qubit 2.0 Fluorometer (Life Technologies) and fragment length and DNA quality was assessed using the Agilent TapeStation 2200 (Agilent Technologies).

Genomic DNA (~11 μg in 100 μL) was fragmented using a Covaris g-TUBE (Covaris, UK) centrifuged at 3380 × *g* for 90 s × 2 to achieve a fragment distribution with a peak at ~16 kb. The sequencing library was prepared using Oxford Nanopore Technologies Nanopore Sequencing Kit SQK-NSK007 (R9 version) according to the manufacturer’s protocol (16 May 2016 version) starting at the End-prep step with ~2.5 ng of DNA. Half (12 μL) of the library was loaded onto a FLO-Min104 (R9 version) flow cell and sequenced for ~22 hours using the script: MinKNOW NC_48hr_Sequencing_Run_FLO-Min104.py. The flow cell was re-started after ~7 hours. The remaining 12 μL of the library was loaded after re-starting the flow cell at 22 hours. Sequencing was then run for a further 43 hours. Base-calling was performed using Metrichor Desktop Agent (v1.107, 2D basecalling for SQK-NSK007).

The complete, raw data set comprised 7,044,217 paired-end 301 bp Illumina MiSeq reads and 53,048 QC-passed Nanopore MinION reads. The Nanopore reads were extracted to fastq format using the poRe R package (61). For the Illumina-only assembly, SPAdes v3.6.2 (62) was used with the k-mer flag set to -k 21,33,55,77,99,127. For the Nanopore-only assembly, Canu v1.5 (63) was used with genomesize=7.0m and the –nanopore-raw flag. For the hybrid Illumina/Nanopore assembly, SPAdes v3.8.2 (64) was used, supplied with both datasets and with the --careful and –nanopore flags. Contigs with low sequence coverage were removed from the hybrid assembly. All assembly tasks were conducted using 16 CPUs on a 256 GB compute node within the Norwich Bioscience Institutes (NBI) High Performance Computing (HPC) cluster. Genome assembly statistics are reported in Table S1. The hybrid assembly genome sequence was annotated using Prokka (65), which implements Prodigal (66) as an *orf* calling tool.

### Cloning the *S. cinnamoneus bcm* gene cluster

The DNA region containing the *bcm* gene cluster was PCR amplified from *S. cinnamoneus* gDNA using primers pIJ-bcm_start and pIJ-bcm_end with Herculase II Fusion DNA polymerase (Agilent). The resulting 6981 bp fragment was gel purified and inserted via Gibson assembly (29) into pIJ10257 (a ΦBT1 integrative and hygromycin resistant vector (28)) linearized with NdeI and PacI to generate plasmid pIJ-BCM. To verify that the cluster sequence in this construct was correct, the plasmid was Sanger sequenced using primers BCM_seq_1 to BCM_seq_8. All other DNA isolation and manipulation techniques were performed according to standard procedures (67).

### Genetic manipulation of *Streptomyces* and heterologous expression of the *bcm* cluster

Methylation-deficient *E. coli* ET12567 carrying the helper plasmid pUZ8002 (68) was transformed with pIJ-BCM by electroporation. This was employed as the donor strain in an intergeneric conjugation with *S. coelicolor* M1146 and M1152 (31), which was performed following standard protocols (59). Exconjugants were screened by colony PCR with primers bcm-cdps_chk_fw and bcm-cdps_chk_rv to confirm plasmid integration. Control strains containing empty pIJ10257 were also generated using the same methodology.

### Cloning and expression of the *P. aeruginosa bcm* gene cluster

Genomic DNA of *P. aeruginosa* SCV20265 was obtained using the FastDNA SPIN Kit for Soil (MP Biomedicals). The DNA region containing genes *bcmA* to *bcmT* preceded by their own native promoter was PCR amplified using primers pJH-BCMclp_start and pJH-BCMcl_end with Herculase II Fusion DNA polymerase (Agilent). The resulting 8604 bp fragment was gel purified and inserted via Gibson assembly (29) into pJH10TS (a derivative of the broad-host-range IncQ expression vector pJH10 carrying the synthetic Tac promoter(39, 40)) linearized with NdeI and XbaI to generate expression plasmid pJH-BCMclp-PA. This plasmid was verified by Sanger sequencing with primers BCM_PA_seq_1 to BCM_PA_seq_9 and introduced into *P. fluorescens* SBW25 via electroporation of freshly made competent cells, which were prepared as follows: two 1 mL aliquots of an overnight culture of *P. fluorescens* were centrifuged at 11,000 × *g* for 1 min, and the pellets washed three times with 1 mL of HEPES buffer each, centrifuging at 11,000 × *g* for 1 min in every wash. The two pellets were then merged and resuspended in 100 μL HEPES buffer and 2 μL of plasmid prep were added to the cell suspension, which was electroporated applying 2500 V. After electroporation, the suspension was transferred to 1 mL of fresh LB and incubated with shaking at 28 °C for 1 hour after which 100 μL of the mixture were plated onto an LB plate containing 12.5 μg/mL tetracycline. As a negative control, the empty vector pJH10TS (40) was also transformed into *P. fluorescens* SBW25. In order to verify the presence and sequence accuracy of the construct in *P. fluorescens*, colony PCR was carried out with transformants using primers pJH_chk_fw and pJH_chk_rv. For the positive clones selected for downstream work, pJH-BCMclp-PA was recovered and sequenced with primers BCM_PA_seq_1 to BCM_PA_seq_9.

### Production and LC-MS analysis of BCM

30 μL of a concentrated stock of *S. cinnamoneus* spores were used to inoculate 10 mL AIZ medium in 50 mL flasks, which were incubated at 28 °C with shaking at 250 rpm for 3 days. 500 μL of this seed culture were used to inoculate 7 mL CIN medium in 50 mL Falcon tubes covered with foam bungs. These production cultures were incubated at 28 °C with shaking at 250 rpm for 4 days. The same procedure was used for *S. coelicolor* M1146-pIJ-BCM and M1152-pIJ-BCM. For production in *P. fluorescens*, 20 μL of cell stocks were used to inoculate 10 mL SCFM in 30 mL universal polystyrene tubes. These cultures were grown overnight at 28 °C with shaking at 250 rpm, with the screw caps slightly loose to allow aeration, and 400 μL aliquots were used to inoculate 10 mL BCMM in 50 mL Falcon tubes covered with foam bungs. Production cultures were incubated for 12-16 h at 28 °C with shaking at 250 rpm.

For the analysis of BCM production, 1 mL production culture samples were centrifuged at 18,000 × *g* for 5 minutes. 5 μL of these samples were analysed by LC-MS using a Luna Omega 1.6 μm Polar C18 column (50 mm × 2.1 mm, 100 Å, Phenomenex) connected to a Shimadzu Nexera X2 UHPLC eluting with a linear gradient of 0 to 35% methanol in water + 0.1% formic acid over 6 minutes, with a flow-rate of 0.5 mL/min. MS data was obtained using a Shimadzu ion-trap time-of-flight (IT-TOF) mass spectrometer coupled to the UHPLC and analysed using LabSolutions software (Shimadzu). MS data was collected in positive mode over a 200-2000 *m/z* range, with an ion accumulation window of 10 ms and automatic sensitivity control of 70% of the base peak. The curved desolvation line (CDL) temperature was 250 °C and the heat block temperature was 300 °C. MS^2^ data was collected between *m/z* 90 and 2000 in a data-dependent manner for parent ions between *m/z* 200 and 1500, using collision-induced dissociation energy of 50% and a precursor ion width of 3 Da. The instrument was calibrated using sodium trifluoroacetate cluster ions prior to every run.

Additional LC-MS analysis was carried out using a Waters Xevo TQ-S Tandem LC-MS fitted with the aforementioned column and employing the same chromatographic method, but injecting 1 μL sample. A multiple reaction monitoring (MRM) method for BCM identification and quantification was configured with Intellistart software (Waters) using pure BCM as a standard. The following transitions were monitored over a dwell time of 0.01 s each (collision energies applied in each case are listed in brackets): for the parent ion with *m/z* 285.11 [M-H_2_O+H]+ : 211.04 (16 V), 193.28 (20 V), 108.13 (28 V) and 81.93 (34 V). For the parent ion with *m/z* 325.10 [M+Na]+: 307.07 (16 V), 251.07 (16 V), 233.18 (20 V) and 215.96 (22 V). Data was acquired in positive electrospray mode with a capillary voltage of 3.9 kV, desolvation temperature of 500 °C and gas flow of 900 L/h, cone gas flow of 150 L/h and nebuliser set to 7.0 bar. LC-MS data were analysed using MassLynx software and the quantification tool QuanLynx (Waters). Xevo MS peak areas were used to determine BCM yields in comparison to a BCM standard.

For the accurate mass measurement of the BCM-like compounds high-resolution mass spectra were acquired on a Synapt G2-Si mass spectrometer (Waters) operated in positive mode with a scan time of 0.5 s in the mass range of *m/z* 50 to 600. 5 μL samples were injected onto a Luna Omega 1.6 μm Polar C18 column (50 mm × 2.1 mm, 100 Å, Phenomenex) and eluted with a linear gradient of 1 to 40% acetonitrile in water + 0.1% formic acid over 7 minutes. Synapt G2-Si MS data were collected with the following parameters: capillary voltage = 2.5 kV; cone voltage = 40 V; source temperature = 120 °C; desolvation temperature = 350 °C. Leu-enkephalin peptide was used to generate a dual lock-mass calibration with *m/z* = 278.1135 and *m/z* = 556.2766 measured every 30 s during the run.

### Isolation and characterization of BCM from *Pseudomonas*

4 × 2L flasks containing 500 mL of BCMM were each inoculated with 20 mL of SBW25-pJH-BCMclp-PA SCFM seed culture grown overnight. After 20 h fermentation at 28 °C with shaking at 250 rpm, the culture broth (approx. 2 L) was separated from the cells by centrifugation to yield a cell-free supernatant (ca. 2 L). The supernatant was lyophilized and then resuspended in distilled water (0.6 L). This aqueous solution was extracted with ethyl acetate (3 × 0.6 L) and then with 1-butanol (3 × 3 L). The solvent was removed to dryness from each extract to afford an ethyl acetate extract (0.014 g), a butanol extract (0.914 g) and an aqueous extract (7.06 g). LC-MS analysis determined that the target compounds were mainly in the butanol and the aqueous extracts. 0.202 g of the aqueous extract and all of the butanol extract were subjected to solid phase chromatography (SPE) on a C-18 cartridge (DSC-18, 20 mL) using a gradient of H_2_O : MeOH (100:0 to 80:20). Fractions containing BCM were combined and further purified by semi-preparative HPLC (Phenomenex, Luna PFP(2), 250 mm × 10 mm, 5 μm; 2 mL/min, UV detection at 218 nm) using a linear gradient of MeOH/H_2_O from 2 to 35% MeOH over 35 minutes, yielding bicyclomycin (3.3 mg, *t_R_* 30.2 min). 1D and 2D NMR spectra were recorded at a ^1^H resonance frequency of 400 MHz and a ^13^C resonance frequency of 100 MHz using a Bruker Avance 400 MHz NMR spectrometer operated using Topspin 2.0 software. Spectra were calibrated to the residual solvent signals of CD_3_SOCD_3_ with resonances at δ_H_ 2.50 and δ_C_ 39.52. Optical rotations were measured on a PerkinElmer Polarimeter (Model 341) using the sodium D line (589 nm) at 20 °C. Commercial standard BCM: [α]^20^_D_ +42.8° (c 0.454, MeOH). *Pseudomonas* BCM: [α]^20^_D_ +43.5° (c 0.091, MeOH).

### Identification of *bcm* gene clusters in sequenced bacteria

The sequences used for the phylogenetic analyses performed in this work were retrieved as follows. A BLASTP search against the NCBI non-redundant protein sequence database was carried out using the CDPS BcmA from *S. cinnamoneus* as the query, and the accession numbers of the resulting 73 hits were retrieved. These accession numbers were then used as input for Batch Entrez (https://www.ncbi.nlm.nih.gov/sites/batchentrez) to retrieve all the genomic records associated to them in the RefSeq nucleotide database (i.e. genomic sequences containing the protein IDs recovered from BLASTP). This yielded a total of 754 nucleotide records which were then analysed using MultiGeneBlast (69) to ascertain which ones had the complete *bcm* gene cluster. 30 of the 754 sequences were discarded on the basis that they did not contain the *bcm* gene cluster or its sequence was truncated. Analysis of the metadata associated with the remaining records lead to the exclusion of 217 *P. aeruginosa* sequences (accession numbers NZ_LCSU01000019.1 to NZ_LFDI01000014.1, ordered by taxonomic ID) in order to avoid overestimation of the cluster conservation, since they were all isolated from a single patient (70).

An additional 134 *P. aeruginosa* sequences (accession numbers NZ_FRFJ01000027.1 to NZ_FUEJ01000078.1, ordered by taxonomic ID) were also excluded from the analysis, due to a lack of associated metadata that prevented an assessment of the diversity of the sample set. Finally, a sequence from accession number NZ_LLUU01000091.1 was also discarded due to the presence of a stretch of undetermined nucleotides (substituted with Ns) in the *bcm* gene cluster. This resulted in a final dataset of 374 sequences: 372 putative *bcm* gene clusters (Data set S1) plus the gene clusters from *S. cinnamoneus* DSM 41675 and *P. aeruginosa* SCV20265. For the downstream formatting of the dataset sequences, scripts or programs that could be run in parallel to process multiple inputs were run via GNU Parallel (71).

### Phylogenetic analysis of the *bcm* gene cluster

Nucleotide sequences of the 374 dataset clusters were trimmed to span a nucleotide region from 200 bp upstream of the start of *bcmA* to 200 bp downstream of the end of *bcmG* (average length of 7224 bp). Phylogenetic analyses were carried out using MUSCLE and RAxML, which were used through the CIPRES science gateway (72) and T-REX (73), and the trees were visualised and edited using iTOL (74).

The 374 trimmed *bcm* gene cluster sequences were aligned using MUSCLE (75) with the following parameters: muscle -in infile.fasta- seqtype dna -maxiters 2 -maxmb 30000000 -log logfile.txt -verbose -weight1 clustalw -cluster1 upgmb -sueff 0.1 -root1 pseudo -maxtrees 1 -weight2 clustalw -cluster2 upgmb -sueff 0.1 -root2 pseudo -objscore sp -noanchors -phyiout outputi.phy The resulting PHYLIP interleaved output file was then used to generate a maximum likelihood phylogenetic tree using RAxML (76). The program was configured to perform rapid bootstrapping (BS) with up to a maximum 1000 BS replicate searches (or until convergence was reached), followed by a maximum likelihood search to identify the best tree, with the following input parameters: Raxml -T 4 -N autoMRE -n correctorientcluster -s infile.txt -c 25 -m GTRCAT -p 12345 -k -f a -x 12345

During the phylogenetic analysis with RAxML, 225 sequences were found to be absolutely identical, and were subsequently removed to allow for a streamlined analysis of cluster phylogeny. After the analysis, sequence with accession number NZ_LLQO01000184.1 was also found to be truncated and was eliminated from the phylogenetic tree, which contained 148 non-redundant entries.

For the phylogenetic analyses of the 2-OG-depedent dioxygenases, the amino acid sequences of BcmB, BcmC, BcmE, BcmF and BcmG from *S. cinnamoneus, P. aeruginosa* SCV20265 and a strain subset including all representatives from *Streptomyces, Actinokineospora, Williamsia, Burkholderia* and *Tistrella*, as well as two from *Mycobacterium* and seven from *Pseudomonas*, were retrieved, aligned with MUSCLE (with same parameters as before except for -seqtype protein -hydro 5 -hydrofactor 1.2) and a maximum likelihood phylogenetic tree was generated with RAxML using the model -m PROTGAMMABLOSUM62, including protein BP3529 from *Bordetella pertussis* (accession ID P0A3X2.1) as an outgroup.

### Analysis of the genomic context of the *bcm* gene cluster

For all of the sequences containing the *bcm* gene cluster, a 20 kb region around BcmA was retrieved and reannotated using Prokka. A subset of these sequences (all Gram-positive bacteria, plus *Burkholderia, Tistrella* and several *Pseudomonas* strains) were analyzed for conserved domains using CDD at NCBI (77) and mobile genetic elements were identified by manual analysis.

## Accession number

The genome sequence of *S. cinnamoneus* DSM 41675 has been deposited in the GenBank database (https://www.ncbi.nlm.nih.gov/GenBank/) with the BioProject ID PRJNA423036.

## ACKNOWLEDGEMENTS

This work was supported by Medical Research Council Newton Fund grant MR/P007570/1 (A.W.T, N.M.V and R.L), a Royal Society University Research Fellowship (A.W.T), Biotechnology and Biological Sciences Research Council Institute Strategic Programme Grants BB/J004561/1 and BB/P012523/1 to the John Innes Centre (A.W.T. and N.M.V), and the NBI Computing infrastructure for Science group through use of its HPC cluster. We would like to thank Susanne Häussler (Helmholtz Centre for Infection Research, Germany) for providing *P. aeruginosa* SCV20265, Jacques Corbeil (CRCHU de Québec-Université Laval, Canada) for providing the Kos collection phylogenetic tree and Barrie Wilkinson (John Innes Centre, UK) for access to MinION equipment and for providing pJH10TS.

